# A staged approach using ants as test organism in ecotoxicology

**DOI:** 10.1101/2025.03.04.641392

**Authors:** Marius Pohl, Mathias Otto, Udo Hommen, Sebastian Eilebrecht, Christoph Schäfers, Benedikt Ringbeck, Bernd Göckener, Jürgen Gadau

## Abstract

Although ants (Hymenoptera: Formicidae) are ecological key components, they are currently not included in ecotoxicity testing. Based on their social organization, ecology and exposure routes to pesticides, we propose a multilevel testing scheme for ants starting from isolated workers (level-1), workers and brood (level-2) and founding queens or entire colonies (level-3). To evaluate feasibility, we tested three ant species (*Camponotus maculatus*, *Crematogaster sp*., *Lasius niger*) using the neonicotinoid imidacloprid as a test substance. Ants were orally exposed through liquid food.

At level-1, worker survival showed a clear concentration response and allowed to estimate LC_50_ with sufficiently narrow confidence intervals. In level-2, worker mortality and sublethal effects negatively affected larval survival and development, e.g. occurrence of naked pupae, with a lower NOEC for larvae (<0.05 mg/L feeding solution) than workers (1.7 mg/L). At level-3, the test substance significantly reduced the reproductive output of newly mated *L. niger* queens over 21 days after a single exposure to 0.5 mg/L.

Our results confirm that ants can be easily handled and tested in the laboratory, making them a valuable extension for ecotoxicity testing. Given their ecological relevance, easy culture in laboratory settings and promising first results, we recommend further research using our new testing scheme, e.g. testing other stressors, species and endpoints. Establishing a robust test protocol for ants would provide coverage of a new and ecologically important group of non-target insects, which contributes to a broader protection of biodiversity and ecosystems.

**Highlights:**

1. Ants play key roles in many ecosystems but are not covered in ecotoxicity testing.
2. We propose a testing scheme for ants with 3 different levels of complexity.
3. Ants can be cultured and tested cost- and space-efficient in the laboratory.
4. Three ant species were tested using imidacloprid in parts of our testing scheme.
5. Lethal and sublethal endpoints can be linked to individual and colony fitness.

**Graphical abstract:** 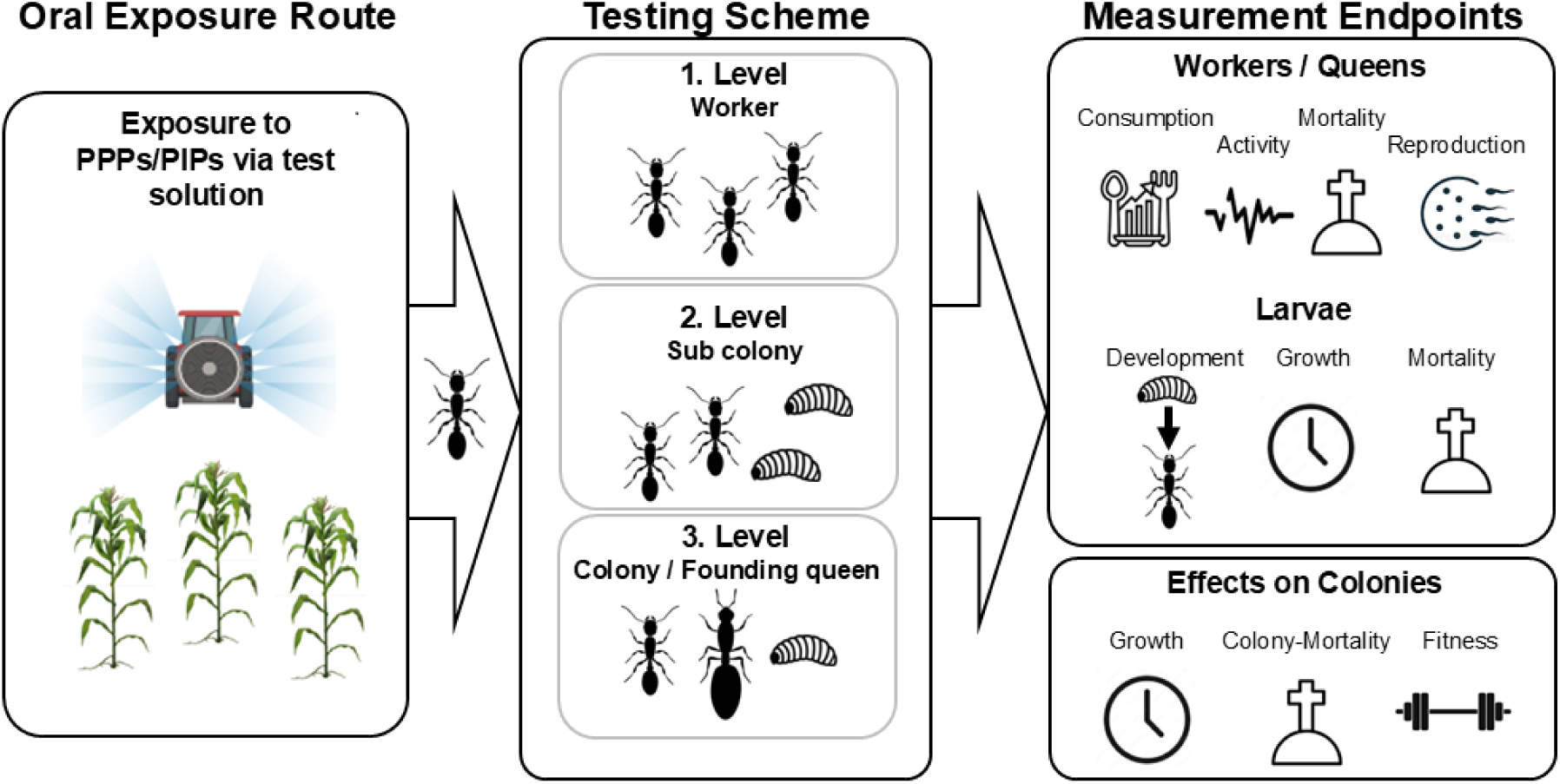

## Introduction

Biodiversity loss is progressing at an alarming rate across all taxa, with particularly insects suffering a significant decline (Hallmann et al., 2017; van Strien et al., 2019). This biodiversity loss is attributed to multiple factors, including habitat destruction, climate change, invasive species, as well as the globally extensive and widespread use of pesticides (Brühl et al., 2021; Habel et al., 2019; Hallmann et al., 2017; Liess et al., 2021; Raven and Wagner, 2021; Ruiz et al., 1997; Vitousek et al., 1958; Wood and Goulson, 2017).

Pesticides, and a range of genetically modified organisms (GMOs) are designed to target pest organisms like insects, weeds or fungi (Kudsk and Streibig, 2003; Morton and Staub, 2008), but can inadvertently affect non-target organisms (NTOs) with severe consequences for food webs, ecosystem services and overall biodiversity (Beaumelle et al., 2023; Brühl and Zaller, 2019; Fritsch et al., 2024; Sánchez-Bayo, 2021). Pesticide application does not only affect cultivated land but can also lead to exposure of natural landscapes and protected areas, resulting in adverse consequences for the corresponding ecosystems (Brühl et al., 2024, 2021). Consequently, protecting biodiversity and addressing pesticides’ ecological risks on NTOs is essential not only for biodiversity conservation but also for maintaining ecosystem services such as pollination, nutrient cycling, soil ventilation and improvement, pest control as well as securing a major food source for various taxa (de Carvalho et al., 2020; Schowalter et al., 2018). While toxicity of pesticides decreased over time for mammals, this is not the case for arthropods (Schulz et al., 2021).

In the EU, the environmental risk assessment of plant protection products (PPPs) is prescribed in Regulation (EC) No 1107/2009 and the one for GMOs in EU Directive 2001/18/EC. Both assessments use a defined range of NTOs, including vertebrate and invertebrate species. Test species are selected for practicability reasons and their function to act as surrogate species which are sensitive to pollutants and relate to ecological functions and/or ecosystem services (e.g. Devos et al., 2016; Romeis et al., 2013). However, the actual protection of the ecosystem also depends on the degree by which set of test organisms represents the diversity of NTOs. This prompts the European Food Safety Authority (EFSA) to emphasize the need of expanding ecotoxicity testing to assess the impacts of pesticides more effectively on biodiversity and ecosystems (EFSA PPR Panel, 2015).

Ants (Hymenoptera: Formicidae) are of high ecological importance, globally distributed and exhibit unique exposure pathways (Del Toro et al., 2012; Hölldobler and Wilson, 1990; Pohl et al., 2024). They are eusocial organisms, exhibiting overlapping generations, shared brood care and reproductive division of labour (Hölldobler and Wilson, 1990) and also feature distinct food ecologies and colony intern food transport processes (trophallaxis) (Meurville and LeBoeuf, 2021). Despite their vital function for ecosystem services, like pollination, soil ventilation, seed dispersal or pest control (Del Toro et al., 2012), ants have been overlooked in regulatory ecotoxicity testing for both PPPs and GMOs. Also, ants exhibit exposure characteristics which are currently not covered in pesticide testing (Pohl et al., 2024). There are only few studies addressing the toxicity of PPPs on ants, with most focusing on ants as target organisms (Bollazzi et al., 2014; Buczkowski, 2016; Wang et al., 2015). Research on ants as a group of NTOs is scarce. However, these already demonstrated sensitivity and adverse effects on survival, reproduction, development and also on behavior (Barbieri et al., 2013; Kwon, 2010; Leponiemi et al., 2022; Rust et al., 2004; Sappington, 2018; Schläppi et al., 2021, 2020; Thiel and Köhler, 2016). Combining all the mentioned characteristics, we see a high relevance in ants being a potential new NTO for future ecotoxicity testing.

Here we propose an approach of testing ants at different levels, based on the social structure and the transfer of toxicants within the colony. We provide an overview over measurement endpoints realized at the different levels and relate them to colony fitness as assessment endpoint. Based on our testing scheme we evaluated all three proposed levels with three different ant species (*Camponotus maculatus*, *Crematogaster sp*., *Lasius niger*) and imidacloprid, an insecticide with known effects on other hymenopteran species (Soares et al., 2015)

### Establishing a concept for testing ants in ecotoxicology

To include ants in ecotoxicity testing, it requires considering aspects of ant biology, exposure characteristics and the need for efficient handling under laboratory conditions. Ultimately, ecotoxicity tests used in environmental risk assessment of chemicals rely on the concept that the tested endpoints are representative for the most sensitive ecosystem structures. As the overall protection goal in our study relates to biodiversity, changes in fitness of an ant colony is supposed to be particularly sensitive and thus can serve as assessment endpoint (see EFSA, 2016). The concept we present addresses colony fitness by testing several levels of organisation. Colony fitness is defined as colony survival and the overall reproductive success and how well genes are propagated to future generations. In comparison to solitary organisms, colony fitness represents the collective performance and reproductive success of the entire social structure (Hamilton, 1964; Hölldobler and Wilson, 1990).

The concept is guided by considering the social structure of colonies as well as exposure pathways (Pohl et al., 2024). The complex organization of an ant colony includes the division of labour, mono- or polymorphic worker castes and the exclusive reproduction by the queen caste. Division of labour in ants is often connected to age-polyphenisms: Younger workers, caring for brood and larvae remain inside the nest. Soldiers usually stay close to the nest guarding the colony and food sources, but they do not actively forage. Foragers feeding the whole colony, although they only make up a small proportion of the worker force (Howard and Tschinkel, 1980; Kloft et al., 1965; Kloft and Hölldobler, 1964). Food transfer between workers, i.e. trophallaxis (the sharing of liquid food) is an important mechanism distributing food within colonies (Kloft and Hölldobler, 1964; Meurville and LeBoeuf, 2021). In the field, foragers may not only be exposed to pesticides by overspray or contact, but also by uptake of contaminated prey or liquid food such as honey dew (Nelson and Mooney, 2022). Oral exposure of workers is the most important exposure route for ant colonies (Pohl et al., 2024) and trophallaxis is the mechanism by which toxicants can be transferred to other workers, brood, queens and alates. Oral exposure can result in chronic exposure at sublethal concentrations.

While the loss of a small portion of the worker caste may be compensated by the colony within a short period of time, the impact on colony fitness becomes more significant when larvae are also affected. Hence, there is an extinction vortex for colonies that loose workers at a constant rate but cannot replace them because larvae are also affected. Additionally, since the ratio of foragers and larvae is usually kept constant, a loss of workers leads to an active reduction of brood size (e.g. by feeding larvae or eggs to other larvae) which further prevents sufficient replenishment of the workforce (Kwapich and Tschinkel, 2013). This eventually leads to a reduction in colony size and eventually to colony death. In any case, the greatest direct impact on colony survival and fitness occurs when the reproductive caste is harmed. If the queen dies and can’t be replaced, the colony ultimately collapses.

Below we explain our rationale and potential measurement endpoints for each of the proposed test levels (see Fig. 1).

**Figure 1:**
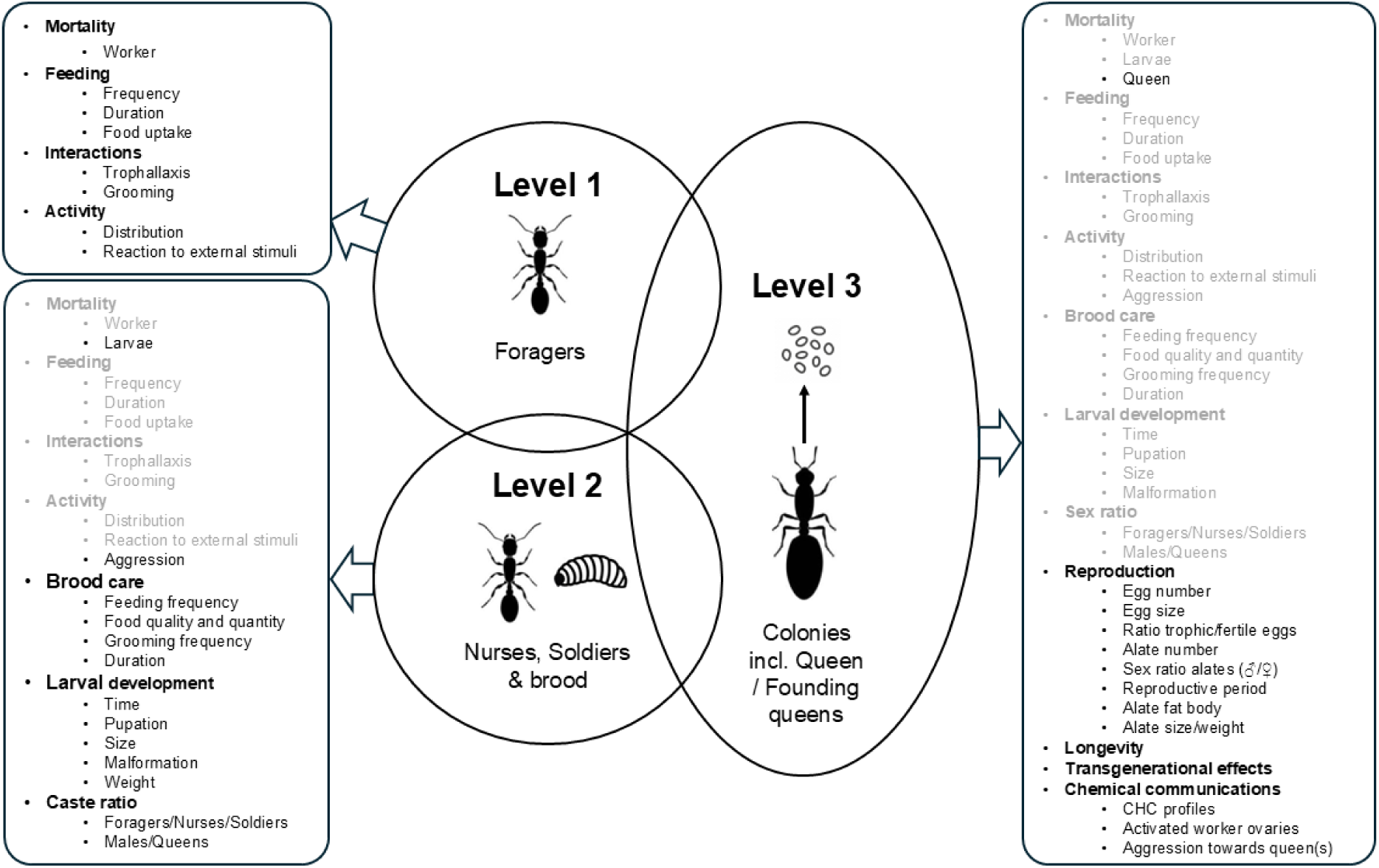
Measurement endpoints for each proposed levels of our testing scheme. Gray points can be tested at a lower or multiple levels. The overlapping individual circles illustrate the staged approach, showing how level-1 and level-2 can be integrated into level-3.

### Level-1 – Workers

Workers specialize in their providing tasks for the colony, e.g. foragers, soldiers, or nurses. This division of labour affects the routes and intensity of exposure to pollutants. Nurses stay inside the nest and thus are only exposed indirectly to toxicants via trophallaxis (Meurville and LeBoeuf, 2021). Soldiers usually stay close to the nest guarding the colony and food sources, but they do not actively forage. In contrast, foragers have the largest activity range collecting liquid food and are the first to encounter any environmentally applied substances. Foraging workers are ideal for assessing the first and direct effects of pesticide because they are responsible for transporting liquid food and prey items back to the colony. Foragers thus face the highest initial exposure regardless of whether topical exposure or exposure via food is considered. As the primary contributors to ecosystem services and caretakers for the brood and queen(s), foragers are ideal test organisms for level-1 studies, as they provide insight into the most direct effects. Additionally, their active foraging behavior allows for testing endpoints such as the acceptance or avoidance of food sources, such as sugary solutions, which can be combined with one or more test substances at varying concentrations.

### Level-2 – Sub-colonies with workers and brood

For the second level, sub-colonies consisting of workers and brood, can be established with the primary aim to assess also the effects of contaminated food, delivered by the workers, on larval survival and development. In the laboratory, multiple small sub-colonies can be established without permanent harm for the initial single queen-right colony. Compared to level-1 effects become more complex, as they may involve different castes and developmental stages (foragers, soldiers, nurses, and larvae). Depending on the mode of action of the test substance, early stages, such as larvae, may be more sensitive than adults (Chen et al., 2022; Lang et al., 2019; Paula et al., 2014). In ants the effects on larvae may be caused either through exposure to contaminated diet via trophallaxis or through starvation due to behavioural changes or death of workers. Diet can be contaminated sugar solution or contaminated prey (e.g. pest organisms which have fed on insecticidal GMPs). Thus, with respect to the exposure of the colony, the test is more complex than the level-1 test restricted to workers.

### Level-3 – Founding queens or whole colonies

Several options exist for experiments in level-3. Here, we suggest two approaches which are related in a more direct way to assess colony fitness than level-1 and level-2. The first one is using entire colonies, including queens. All reproduction in ant colonies relies on the queen caste which requires high volume and high-quality nutrition to constantly produce eggs. Furthermore, ant colonies require little space and can be easily handled in the laboratory, where testing whole colonies is possible and allows a range of direct and indirect endpoints to be measured (Fig. 1). Toxicants may directly harm the queen caste or manifest in transgenerational effects on larvae if toxicants are transferred to eggs. However, in many cases, a colony will collapse if the queen caste is killed.

While testing level-1 and level-2 can be accomplished with a single colony from which sub-colonies of foragers, nurses or larvae can be used, several colonies are needed testing level-3. Thus, the effort in culturing is high. Therefore, we also propose a less complex alternative for level-3, which allows to test effects on survival and reproduction of queens. This approach does not need entire colonies but only founding queens, which can be available in larger numbers than entire colonies.

The second approach would be testing individual founding queens right after the nuptial flight, which can be seen as a proxy for the reproductive caste of a colony. The advantage of testing founding queens would be that no entire colony would be sacrificed for testing, as large, thriving ant colonies need extended time to grow. Number and survival of alates also are meaningful for colony dispersal and included in assessing the overall fitness of colonies.

### Measurement endpoints and their significance for colony fitness

Proposed levels and respective endpoints all relate to changes in colony fitness, which we consider as the most relevant assessment endpoint (EFSA, 2016). As the levels progress, the complexity of the experimental design increases, considering the biology and social structure of ants. Direct and indirect effects on different castes can influence colony fitness in different ways: For instance, the loss of foragers will impact the available food and thus the available energy for the colony. The loss of nurses can impact the feeding frequency towards larvae and may even cause a delay in development.

The loss of soldiers/majors (subcaste, e.g. in *Camponotus* spp. or *Pheidole* spp.) will affect division of labour and the protection of a colony and might lead to an increased number of intruders into the colony (Huang, 2010; Lillico-Ouachour and Abouheif, 2017; Lumsden and Hölldobler, 1983). A significant reduction of nurses will often lead to an active reduction of brood by workers, because there seems to be a standard ratio of worker to brood (Kwapich and Tschinkel, 2013)

The loss of alates, i.e. winged unmated queens and males, directly impacts reproductive success of a colony and its fitness. Diminishing brood numbers or extended developmental times of larvae can negatively affect colony growth. Particularly in the case of European ant species, colony size at the beginning of winter is critical to survive hibernation (Shiroto et al., 2011). The queen caste solely consists of mated female(s) in a colony to produce workers and any negative effect on the queen caste will strongly affect the colony and colonýs fitness. In many ant species, colonies do not accept a newly mated queen if the original queen(s) dies, leading to a collapse of the entire colony and a colony fitness of zero. The queen also controls workers by pheromones and cuticular hydrocarbons (CHCs) and provides the colony with a unique smell. This smell is crucial for communication between colony members and interaction with members from other colonies. An overview of potential mechanisms affecting colony fitness is provided in Figure 1.

In social insects, like ants, a colony is the equivalent of an individual in a solitary species (Boomsma and Gawne, 2018; Hölldobler and Wilson, 2009; Reeve and Hölldobler, 2007; Wheeler, 1911). Hence, colony survival is the ultimate measurement endpoint. However, everything that negatively effects colony growth and survival of individual castes or developmental stages can have detrimental effects on colony survival and is therefore an important measurement endpoint. Social insect colonies are buffered by workforce and the fact that the majority of individuals are in a protected nest environment. Therefore, social insect colonies will be less affected by chemical insecticides over a short period in the same way as solitary non-target organisms. Thus, indirect exposure pathways and sublethal effects as measurement endpoints can be very important to evaluate the effects of a toxicant for social insects.

### Selecting ant species

Many ant species are already maintained under laboratory conditions for years to investigate various topics, e.g. genetics, chemical communication, behavior etc. This already gives possibilities to use a big variety of species to use them in ecotoxicity testing. As laboratory settings are usually simple, space and work efficient, standardized keeping conditions and repeatability are the basis for ecotoxicological testing (Andow et al., 2004).

Because we aimed to establish a standardized exposure concept of oral uptake using honey water mixed with the pollutant, we considered only genera or species with comparable exposure pathways and intracolonial food distribution mechanisms. We also only selected omnivorous ants, which consume a variety of liquid and solid food sources, i.e. were known to consume carbohydrates in form of sugary solutions. In addition, we only chose species that use stomodeal trophallaxis (Meurville and LeBoeuf, 2021) to ensure that the ingested food solutions are distributed from one individual to another. That left us with representatives of the two most common and dominant European ant subfamilies, Formicinae and Myrmecinae (Seifert, 2007; Ward, 2007). Both are represented in temperate and tropical regions (Hölldobler and Wilson, 1990), allowing for the potential inclusion of more subfamilies with similar exposure model in future studies. Furthermore, we aimed to choose test species of different sizes because substantial morphological differences could account for size-related sensitivity The three species chosen were as follows:

***Lasius niger*** *(LINNAEUS* 1758; Black Garden ant) is a widespread and prevalent formicine species in Europe and Asia and commonly distributed in agricultural fields and man-made environments (Seifert, 2020). They build subterranean colonies that exhibit colony sizes between 4000 to 7000 individuals and only include a monotypic worker caste with sizes varying from 3,5-4,5 mm. *L. niger* is omnivorous and favours honey dew produced by hemipterans as carbohydrate source (Detrain and Prieur, 2014). Colonies can exhibit facultative pleometrosis, but become secondary monogynous after the emerging of the first workers (Sommer and Hölldobler, 1995). As a species adapted to the climate of the northern hemisphere, the development of *L. niger* colonies requires a prolonged hibernation period during winter months after which they need to recover for several month to reach full colony size (Tauber et al., 1986). *L. niger* disperses through nuptial flights of alate sexuals, which can easily be collected in spring. As one of the most common ant species in European landscape, we selected *L. niger* as one of our test species due to its accessibility and high potential to be exposed under European conditions in farmed land.

***Camponotus maculatus*** *(FABRICIUS* 1782; Carpenter ant) is a highly diverse formicine species, including various subspecies, widely distributed in Africa, the Middle East and partly in Asia and Australia. *C. maculatus* is a ground nesting ant species (Taylor et al., 2016) that exhibit colony sizes of several tens of thousands of workers and is a rather tropical, i.e. does not hibernate. It represents a large, dominant ant species with a highly polymorphic worker caste (minors, intermedia and majors), ranging in size from 8-16 mm. Their diet is generally omnivore and they extensively feed on nectar and honey dew (Hölldobler and Wilson, 1990).

*Camponotus* is a diverse and globally distributed genus with mostly simple colony structures (monogyny) and *C. maculatus* was chosen for its easy maintenance, easy accessibility of queens (annual nuptial flights), big colonies and realistic exposure to agricultural pesticides. Despite its tropical distribution, closely related species are also distributed in European (e.g. *C. herculeanus* and *C. ligniperdus*) areas and *C. maculatus* was especially selected due to its high growth rate and availability for ecotoxicity testing all year around.

***Crematogaster sp*.** (*LUND* 1831; Acrobat ant), is globally distributed and highly diverse myrmicine genus, with more than 400 species (Blaimer, 2012). Colony size ranges from few hundred up to several thousand individuals (Feldhaar et al., 2003; Heil et al., 2001). Worker size can vary between species but rarely exceeds 5 mm. Caste determination and social structure are diverse in this ant genus, including mono- and polymorphic worker caste (minor and majors) and mono- and polygynous colony structures (Gotoh et al., 2017). *Crematogaster* is an ecological dominant and omnivorous genus (Richard et al., 2001) with nests can be found both arboreal and subterranean.

We selected a yet unidentified *Crematogaster* as a representative of a different subfamily (Myrmecinae). *Crematogaster* has a very high reproductive potential and is easily controlled and reared in the laboratory.

## Material & Methods

### Collection and maintenance of ant colonies

*Camponotus maculatus* and *Crematogaster sp*. founding queens were collected in the Comoé National Park, Ivory Coast in 2019 using a light trap during nuptial flights. The necessary collection permits for the Ivorian ants were provided by the Directeur Général of the Office Ivoirian des Parcs et Réserves (OIPR), Côte d’Ivoire (N°018/MINEDD/OIPR/DZ). Colonies were since then maintained in the facilities of the University Münster. *Lasius niger* founding queens were collected in Münster, Germany in July 2021 during nuptial flights.

Stock colonies were kept in IKEA Samla boxes (article number: 694.408.36 for *C. maculatus*) or rectangular plastic boxes (20 cm (width) × 9 cm (height) × 20 cm, Dino AG, Berlin, Germany for *Crematogaster sp*. and 10 cm (width) × 9 cm (height) × 20 cm, Dino AG, Berlin, Germany for *L. niger*). The base of the boxes was covered with plaster for *Crematogaster sp.* and *C. maculatus*. Nest boxes include 1-3 cavities, depending on colony size, which were covered with red foil, to simulate designated nesting chambers. Box bases for *L. niger* were covered with concrete and an artificial chamber-system, including three cavities, all covered with red foil. Lids had four ventilation holes (IKEA Samla Ø = 6 cm, rectangular Box Ø = 4 cm) covered with metal mesh. *C. maculatus* and *Crematogaster* were kept at 30°C day and 25°C night temperature, while *L. niger* was kept constant at 25°C for 9 months. *L. niger* colonies were annually hibernated at 4°C for the remaining 3 months. The day and night cycles were adjusted to 12-hour light and 12-hour darkness for *C. maculatus* and *Crematogaster*. *L. niger* colonies were kept in constant darkness. The humidity for all three species was set to 50%. Stock colonies were fed *ad libitum* with cockroaches (*Blaptica dubia*) and pure honey twice a week. Water supply in form of water filled tubes, clogged with cotton was constantly available. *C. maculatus* and *Crematogaster sp.* colonies had at the time of the experiments more than 2000 workers, *L. niger* colony sizes ranged from 50 to 100 workers.

### Test substance and solutions

Exposure was realized via test substance solved in 10%-honey solution. Imidacloprid as test substance was purchased from Sigma Aldrich (Taufkirchen, Germany) as an analytical standard. Imidacloprid test solutions were freshly prepared for each test. As a stock solution, 10 mg of the analytic standard was dissolved in 100 ml of deionized water at 50°C. The stock solution was used to prepare seven subsequent test concentrations using deionized water (0.05, 0.17, 0.5, 1.7, 5, 17 and 50 mg/L). For each test concentration and the negative control consisting of deionized water, 50 mL test solution was prepared containing 10% honey to provide an attractive food source for ants. Each solution was mixed on a Vortex (Genie 2 G560E, Scientific Industries, New York, USA) for approximately 5 minutes until the honey was fully dissolved. The test and stock solutions were stored at 4°C and used for a 10-day exposure experiment. For 21-day exposure experiments, the test solutions were renewed using the same stock solution after 10 days of the exposure period.

### Chemical analyses of test solutions

Nominal concentrations of the imidacloprid solutions were confirmed using one millilitre of freshly prepared test solution, which was separately stored at –20°C until chemical analysis. For chemical analysis, ultra-high-performance liquid chromatography-tandem mass spectrometry (UHPLC-MS/MS) was used, and concentrations of test solutions were determined using isotope dilution analysis with imidaclorid-d_4_ as an internal standard. For the measurements, the test solutions were stabilized with acetonitrile in a ratio of 5+1 (test solution + acetonitrile, v/v) and then diluted with a water/acetonitrile mixture with the same composition to fit into the calibration range of 3 to 10,000 ng/L.

To validate the intended exposure, we measured the imidacloprid concentrations in the sugar solutions with the seven concentrations tested in one of the experiments (Cm1, see Tab. 1). Details of the method are described in the Supplementary Material (SI-1 and 2).

**Table 1:** Overview of our five experimental settings for all three testing levels using three ant species and imidacloprid. For the different test concentrations see Material & Methods (range: 0.05 – 50 mg/L).

| Test design |  |  |  |  |  |  |  |
| --- | --- | --- | --- | --- | --- | --- | --- |
| Level | Experiment | Species | Tested stages | Exposure h/day | Test duration (d) | # test conc. (incl control) | Observations |
| 1 | Cm1 | *Camponotus maculatus* | W* | 24 | 10 | 8 | Survival |
| 1 | Ln1 | *Lasius niger* | W | 1 | 10 | 8 | Survival |
| 1 | Cs1 | *Crematogaster sp.* | W* | 1 | 10 | 8 | Survival |
| 2 | Cm2 | *Camponotus maculatus* | W+L** | 1 | 21 | 8 | Survival, Feeding rate, pupation, pupal weight |
| 3 | Ln3 | *Lasius niger* | Q*** | **** | 21 | 4 | Survival, egg production |

Cm1=level-1 and Cm2=level-2 (*Camponotus maculatus*); Ln1=level-1 and Ln2=level-3 (*Lasius niger*) and Cs1=level-1 (*Crematogaster sp.*); \*Worker; \*\*Worker and Larvae; \*\*\*Founding queen. \*\*\*\*Founding queens were only exposed on the first day with a single dose (10 µL) for each test concentration.

The measured concentrations ranged between 101 – 119 % of the nominal concentrations. Therefore, we could confirm the correct preparation of the feeding solutions hence used the nominal concentrations for characterizing the different exposure levels in the following.

### Testing procedure and endpoints

We conducted a total of five experiments covering our proposed three different levels. Three ant species were used for experiments of level-1, while only one species was used for level-2 and level-3, respectively (see Tab. 1).

Sub-colonies comprising 50 workers, or 50 workers and 20 larvae were kept in plastic boxes (10 cm (width) × 9 cm (height) × 20 cm (depth), Dino AG, Berlin, Germany) with two ventilation holes (Ø =4 cm) in the lid, covered with metal mesh. The base of the boxes was covered with plaster for *C. maculatus* and *Crematogaster* or concrete for *L. niger*, and cavities covered with red foil were added to mimic artificial nesting chambers (SI 2). While *L. niger* and *Crematogaster* exhibit monomorphic worker castes, for *C. maculatus* minor workers randomly chosen from stock colonies were used. For each experiment, six sub-colonies were used as controls and three sub-colonies for each of the seven test concentrations. The workers for each sub-colony originated from a single stock colony, comprising three stock colonies of *C. maculatus* and *Crematogaster sp*., while each sub-colony originated from a separate stock colony for *L. niger*.

The sub-colonies were exposed daily to 750 µL and 200 µL test solution, for *C. maculatus* and *L. niger*/*Crematogaster,* respectively, while a single leg of *Blaptica dubia* were provided as protein source every other day. Only for experiment 1 (Tab. 1, Cm1), sub-colonies were exposed continuously for 24 hours each day. Due to strong evaporation, which led to unquantifiable food intake rates. In all other exposure experiments (Tab. 1), exposure period was adjusted to one hour/day to minimize the evaporation of the test solution, to be able to calculate the amount of test solution taken up by workers. This was still sufficient time for workers to feed. Additionally, we controlled for evaporation by measuring it in parallel to our experiments in an empty nest box. This evaporation control was used to correct the calculation of food intake in the experimental groups for evaporation. For level-1 experiments containing only worker sub-colonies, the exposure was conducted over 10 consecutive days (Cm1, Ln1, Cs1, Fig. 2). The individual daily intake was calculated based on the number of workers exposed each day and the sub-colony’s consumption rate after correcting for evaporation. For the two species, *Lasius niger* and *Crematogaster sp.*, a qualitative confirmation of the test solution uptake and transfer to other individuals via trophallaxis was conducted. Therefore, food dye was added to the test solution colouring the crop and gut of the ants (SI-6).

**Figure 2:**
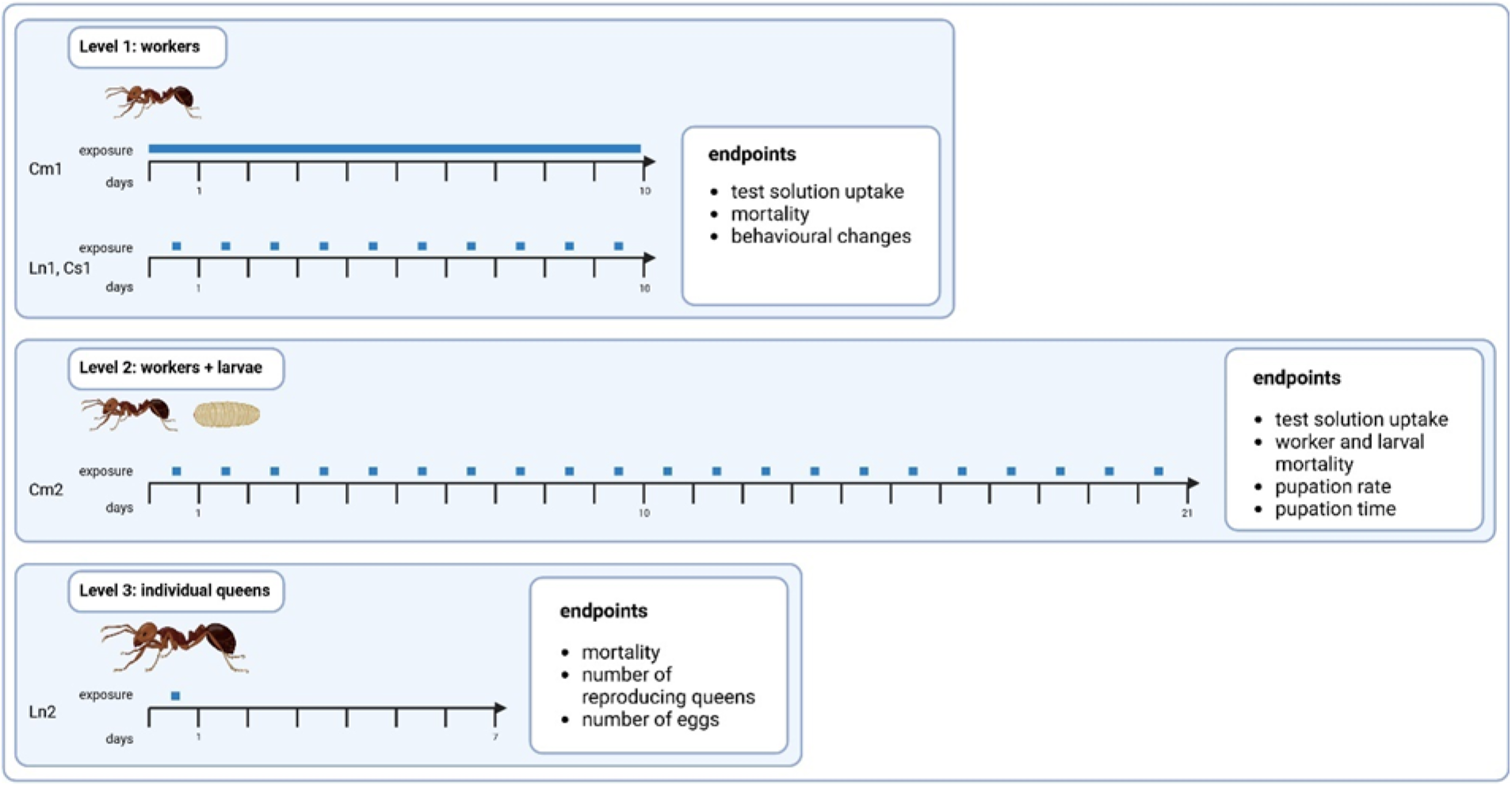
Graphical overview of our experimental design illustrating levels 1–3; exposure characteristics (24 h/day or 1 h/day; single exposure for level-3 or repeated exposure for levels 1 and 2, represented by blue boxes or bars above the time axis in days), and measured endpoints.

In the experimental setup of level-2 (Cm2, Fig. 2), sub-colonies consisting of 50 worker and 20 second-instar larvae were exposed to the test solutions over a 21-day period. During this time, larvae typically grow significantly, enabling the assessment of developmental effects during larval development using this setup. Following the 21-day exposure to the test solutions, sub-colonies were provided with a regular diet of honey and *B. dubia* legs. The experimental sub-colonies were monitored until all larvae had either pupated or perished. Any larvae that were absent, presumably consumed by the workers, were recorded as deceased. We subsequently assessed worker and larval mortality, pupation rate, and pupal weight. Worker mortality was assessed daily, while larval mortality was monitored every other day to reduce handling stress.

To represent level-3, we used founding queens of *L. niger*, collected after the nuptial flights beginning of June 2024, and exposed them individually to three different imidacloprid concentrations (0.5, 5 and 50 mg/L) and to a negative control (n=20). After collection, queens were kept in 4 mL vials (article number: K937.1, Carl Roth GmbH + Co. KG, Karlsruhe, Germany), with a water reservoir and the vial openings sealed with cotton. The queens were monitored for 10 days prior to the experiment and only those that laid eggs were selected for experiments to ensure that only mated and reproducing queens were included. For level-3 (Ln3), the queens were transferred to a new vial and fed with 10 µL of the test solutions via a 0.2 mL pipette tip (Sarstedt AG & Co. KG, Nürnbrecht, Germany) at day 0. Founding queens were exposed once to simulate conditions during the nuptial flight, i.e. when they are exposed to the environment rather than being protected in a soil nest. The founding queens were observed for three weeks and perished queens as well as individually laid eggs were counted afterwards.

### Statistical analyses

NOEC / LOECs for survival or other quantal data were calculated by the step-down Cochran-Armitage test procedure. Effects on feeding rate, day of pupation, body weight, and reproduction were analysed by appropriate multiple tests after testing on normal distribution and variance homogeneity. The replicates (i.e. 6 controls and 3 test groups per replicate) were considered as experimental units. However, for day of pupation and pupae weight in experiment Cm2 and for reproduction per survivor (queen) the data per individual pupae and queen were used, respectively. The significance level was set to 0.05, one-sided.

Survival and other quantal data were analysed by probit analysis to calculate LC_X_ values based on nominal concentrations in the test solutions because the measured concentrations were close to the nominal ones (+/- 20%). In addition, LD_X_ were calculated in the Cm2 experiment, in which the daily feeding rates were determined. Therefore, the mean daily uptake of imidacloprid per ant was calculated from the mean daily feeding rates over time and the nominal concentrations of the test item in the honey solutions. The mean measured wet weight per ant (SI-4 based on measurements of *C. maculatus* minor workers) was used to estimate the mean daily dose per biomass. For non-quantal data like weight or reproduction, a 3-parameter log-logistic model was used to calculate EC_X_ values.

The software ToxRat Professional 3.3 (ToxRat Solutions, Alsdorf, Germany) was used for statistical analysis. Statistical results are summarized in Table 2 and the detailed results are provided in the repository.

**Table 2:** Summary of the statistical results of all experiments. All concentrations are given in mg/L in our food solution (see methods).

| Exp. | Caste | Exposure | Trait | NOEC / LOEC | p-value<br>LOEC | % effect<br>NOEC / LOEC | Test | L(E)C<sub>50</sub><br>[mg/L] | 95 % CI | % control<br>mortality | Model |
| --- | --- | --- | --- | --- | --- | --- | --- | --- | --- | --- | --- |
| Cm1 | Wo | 10 days<br>24 h/day | 10-day survival | 5 / 17 | <0.001 | 39 / 74 | RSCA | 8.35 | 4.82 - 15.5 | 4.3 | probit |
| Ln1 | Wo | 10 days<br>1 h/d | 10-day survival | 0.05 / 0.17 | 0.027 | 0 / 2 | RSCA | 11.0 | 6.30 - 21.9 | 0.0 | probit |
| Cs1 | Wo | 10 days<br>1 h/day | 10-day survival | 0.17 / 0.5 | 0.010 | 9 / 15 | RSCA | 3.97 | 3.27 – 4.84 | 0.0 | probit |
| Cm2 | Wo | 10 days<br>1 h/day | 10-day survival | 0.5 / 1.7 | 0.031 | 5 / 29 | RSCA | 11.2 | 6.77 – 20.9 | 5.3 | probit |
| Cm2 | Wo | 10 days<br>1 h/day | 10-day<br>mean feeding rate | ≥ 50 / > 50 | - | - | Welsh | n. d. | - | - |  |
| Cm2 | La | 10 days<br>1 h/day | 10-day survival <sup>a</sup> | ≥ 50 / > 50 | - | 26.7- | RSCA | n. d. | - | 24 |  |
| Cm2 | La | 10 days<br>1 h/day | 10-day survival<br>up to 5 mg/L | 0.5 / 1.7 | <0.001 | 6.7 / 37 | RSCA | n. d. | - | 24 |  |
| Cm2 | La | 10 days<br>1 h/day | survival<br>until pupation | 0.5 / 1.7 | <0.001 | 70 / 88 <sup>c</sup> | RSCA | 0.77 | 0.61 – 1.02 | 54.2 | probit |
| Cm2 | La | 10 days<br>1 h/day | day of pupation <sup>d</sup> | ≥ 0.5 / > 0.5 | - | +86 / - | Williams | n. d. | - | - |  |
| Cm2 | Pu | 10 days<br>1 h/day | building normal pupa | < 0.05 / 0.05 | 0.007 | 0 / 11 | Chi<sup>2</sup> | n.s. | - | 0 | probit |
| Cm2 | Pu | 10 days<br>1 h/day | body weight <sup>b</sup> | < 0.05 / 0.05 | 0.008 | < 13 / 13 | JT | 18.1. | n.d. | - | 3-parameter<br>logistic |
| Ln3 | Qu | 1 days<br>1x | 21-day survival | 5 / 50 | <0.001 | 15 / 40 | RSCA | 111 | 34.4 – n.d. | 5.0 | probit |
| Ln3 | Qu | 1 days<br>1x | 21-day<br>reproduction | < 0.5 / 5 | n.d. | < 18 / 18 | Williams | 259 | n.d. |  | 3-parameter<br>logistic |

n.d. = not determined, n. s. = no significant regression, RSCA = Step-down Rao-Scott-Cochran-Armitage test procedure, Welsh = Welsh t-test after Bonferroni Holm, Chi<sup>2</sup> 2x2 table test with Bonferroni correction, JT = Step-down Jonckheere-Terpstra, 3 para logistic = 3 parameter logistic cumulative density function
<sup>a</sup> No monotonous concentration response of larval survival until day 10: control mortality = 24.2 % and at 50 mg/L 26.7 % while the highest mortality was found at 5 µg/L (58.3%). No significant effects according to Rao-Scott-Armitage test procedure. <sup>b</sup> Individual body weights were used for analysis, not mean body weight per replicate. <sup>c</sup> Pupation rate of the control was 54%. <sup>d</sup> In 1.7 mg/L only two pupae were found. Thus, effects on day of pupation (increase) were only analysed up to 0.5 mg/L. Note that for practical reasons, the experiment could not start with only larvae of the same stage. <sup>e</sup> LC<sub>50</sub> extrapolated above the highest test concentration of 50 mg/L.

## Results and Discussion

### Test solution uptake

Pesticides and their formulation components, particularly when ingested orally, can be detected by various organisms, resulting in diverse levels of acceptance or even avoidance behaviors of the food source (Frizzi et al., 2020; Rohel et al., 2001). Ensuring test solution acceptance is crucial for accurately assessing a toxicant’s hazardous potential. Without the verification, results could be misleading and/or mitigated. Therefore, the initial step in our testing scheme was to confirm that foragers consumed the test solution. Overall, all species tested readily consumed the imidacloprid - honey water solutions at all tested concentrations without hesitation.

In the level-2 experiment with *C. maculatus* (Cm2), consumption rates of solutions with different imidacloprid concentrations show no clear trend over the lower test concentrations but at the highest test concentration of 50 mg/L, the mean consumption rate was reduced by 58 %, but not statistically significant (Welsh t-test after Bonferroni correction, p = 0.021, α = 0.007). However, at the highest concentration of 50 mg/L, workers experienced significant lethal and sublethal effects, which likely affected their individual intake over time. Because the consumption rate was measured uniformly over 10 days and individuals initially consumed the test solution without avoidance, the immediate sublethal and lethal effects likely explain why the reduction in feeding was not statistically significant (SI-5). A dynamic adjustment of the consumption rate based on actual feeding behavior over time could offer a more precise evaluation of its impact.

Consumed volumes, ingested by *L. niger* and *Crematogaster* workers, were too small to be measurable with sufficient precision. However, observation of feeding behavior, along with similar effects on survival for all three species, indirectly confirmed that all test solutions were ingested by both *L. niger* and *Crematogaster sp*.

### Level-1 – Effects on workers

#### Worker mortality

Imidacloprid is a systemic insecticide and binds irreversibly to the nicotinic-acetylcholine receptors in the insect’s nervous system (Tennekes, 2010). Thus, its effects can accumulate over time and lead to disruption of nerve signalling, convulsions and loss of coordination (Casida and Durkin, 2013; Parkinson and Gray, 2019), ultimately leading to death. Worker mortality over time (SI-7) showed significant concentration response relationships with narrow confidence intervals for all three species (Fig. 3). LC_50_ for survival over 10 days for *C. maculatus* and *L. niger* were very similar (8.35 – 11.03 mg/L), while the LC_50_ for *Crematogaster sp.* was about a factor of two lower (3.97 mg/L). NOECs calculated by the Cochran-Armitage tests were 5, 0.05 and 0.17 mg/L for *C. maculatus*, *L. niger* and *Crematogaster sp.,* respectively. However, the ecotoxicological relevance of the NOECs alone is low because significant mortality levels vary between 2 % (*L. niger*) and 74 % (*C. maculatus*).

**Figure 3:**
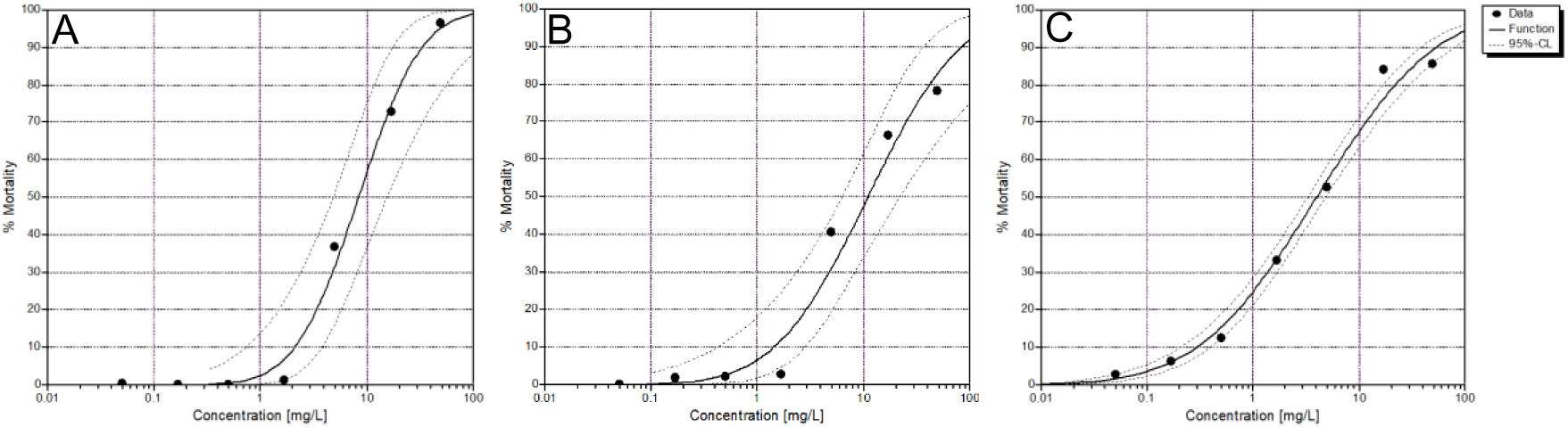
Concentration response curves fitted by probit analysis for survival over 10 days of workers in the Level 1 experiments with three ant species:(**A:** *Camponotus maculatus* (Cm1), **B:** *Lasius niger* (Ln1) and **C:** *Crematogaster* sp. (Cs1)). Exposure is given as nominal imidacloprid concentrations in the food solutions.

LC_50_ values for our three tested ant species were in the same order of magnitude as LC_50_ values based on oral exposure for honey bees (Wong et al., 2018) and stingless bees (Al Naggar et al., 2022; Costa et al., 2015) following oral exposure. However, a direct comparison is challenging, because most bee toxicity tests are conducted with an exposure time of 24-48 h, whereas our LC_50_ values are based on a chronic 10-day exposure. Comparing our data to the effects of imidacloprid on honey bees, it seems that bees are more susceptible to immediate lethal effects. Aljedani (2017) reported mortality in *Apis mellifera* within 4 hours, whereas in *C. maculatus*, lethal effects began to occur after 48-hour.

Delayed sublethal and lethal effects, especially for chronic field realistic exposure, can be considered to have similar or even more severe consequences for social insects than short-term exposure to higher concentrations. As imidacloprid persists over an extended period of time in field environments, the time-cumulative toxicity should be assessed using chronic exposure over a longer time (Rondeau et al., 2014).

### Sublethal effects

Two types of behavioural changes were observed for *C. maculatus* workers with increasing imidacloprid concentrations. First, overall activity increased leading to a more even distribution of workers in the experimental box (SI-3) compared to the control. The NOEC for this behaviour was determined to be 0.17 mg/L. A possible explanation for the increased activity and even distribution may be related to increased foraging for alternative food sources or the known neurological effects of imidacloprid (Casida and Durkin, 2013; Parkinson and Gray, 2019). Bortolotti et al. (2003) showed that imidacloprid can impair the homing behaviour of honey bees and thus could have a similar effect on the orientation system of ants. In future experiments, behavioral video tracking combined with automated analyses could be used to evaluate sublethal effects at the behavioral level, such as feeding frequency (worker – worker, worker – larvae), intracolonial interactions, homing behavior, and overall activity. Currently, various tracking systems for insects exist that allow for continuous monitoring of individual workers over extended periods without disturbing them (Rydhmer et al., 2022).

Secondly, ants appeared to show a reduced response to external stimuli. The videos provided in the data repository show *C. maculatus* workers with compromised locomotion and coordination after being exposed to test solutions with 5 mg/L imidacloprid concentration. Here, we assessed this effect only qualitatively. Behavioral changes on orientation caused by imidacloprid have been also observed in the locust species *Locusta migratoria.* In this species imidacloprid exposure disrupted nerve signalling and caused loss of coordination due to neurotoxic effects (Parkinson and Gray, 2019).

### Summary

In general, worker mortality can be used as a straightforward, reliable, and easily measurable endpoint for all levels of our testing scheme. Furthermore, we consider worker behaviour to be a valuable measurement endpoint for social insects, because behavioral changes can have profound negative effects on essential behavioural traits, such division of labour, foraging or brood care, which all have a negative influence on colony growth and are therefore highly relevant for colony fitness.

### Level-2-Effects on workers and larvae in sub-colonies

#### Effects on workers

Effects on *C. maculatus* worker survival in the level-2 test (Cm2) and behavior are comparable to those observed in level-1. The LC_50_ in Cm2 was slightly higher than in Cm1: 11.2 mg/L (Fig. 4A) compared to 8.4 mg/L (Fig. 3A). The NOEC was lower (0.5 compared to 5 mg/L) but mortality at the NOEC was different with 39 % in Cm1 and 5% in Cm2.

**Figure 4:**
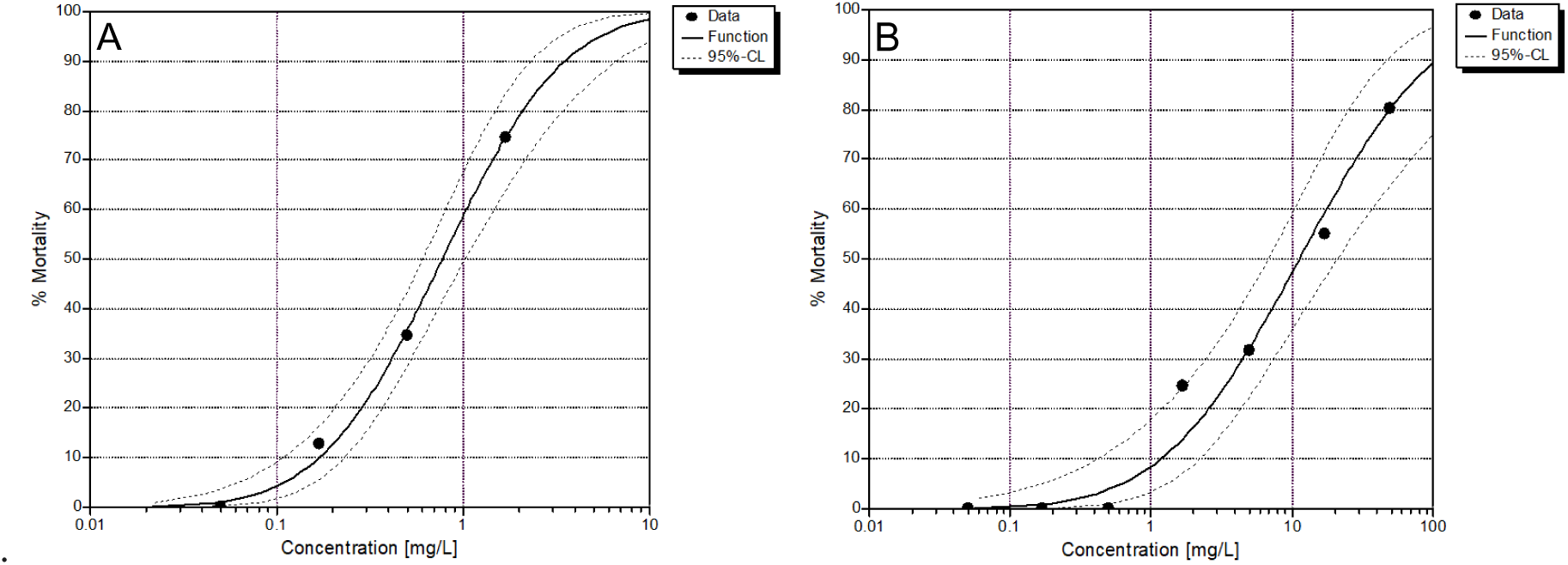
**A:** Mortality of *C. maculatus* workers until day 10 and **B:** Larvae until pupation in the Cm2 experiment. Control mortality was compensated by using Abbott’ s formula.

Since consumption rates of workers were measured continuously in the Cm2 experiment, daily imidacloprid doses per ant could also be estimated. Based on the mean daily uptake rates related to the mean worker body weight (13.12 mg fresh weight, SI-4), The LD_50_ for *C. maculatus* workers was calculated at 10.67 ng/ant over a 10-day exposure period. This value falls within the same order of magnitude as reported LD_50_ values for honey bees, which range from 3.7 to 40.9 ng/bee for an exposure duration of 48–96 hours (Schmuck et al., 2001; Suchail et al., 2001, 2000). However, direct comparison remains challenging, as ants in general are smaller than honey bees and the exposure durations differ. One approach to account for size differences would be to scale the dose per wet weight (g) and day, resulting in an LD_50_ of 0.813 µg/g wet weight/day over a 10-day exposure period (95% CI: 0.544–1.33) for *C. maculatus* workers. Except for honey bees, exposure in standardized ecotoxicological tests is rarely expressed in doses but to a concentration over time. However, the use of doses is preferable if TD/TK models are to be used for calculation of risks.

### Larval mortality and development

Larval mortality until day 10 showed a non-linear pattern, it increased till the second highest concentration (53.8 %, 0.5 mg/L), but decreased again at the highest concentration (26.7%, 50 mg/L) and became indistinguishable from our controls (24.8%). The step-down Rao-Scott-Cochran-Armitage test procedure did not find a significant difference to control, except if the data for the two highest test concentrations are ignored (NOEC = 0.5 mg/L). Thus, LC_X_ calculation was not performed. The observed mortality patterns can be explained by the reliance of ant larvae on brood care provided by workers. Larvae in our tested species are exclusively fed through trophallaxis and are incapable of surviving independently. Furthermore, since larvae always stay in the protected surroundings of the nest, they can only be indirectly exposed to toxicants through feeding contaminated food. Larvae are highly nutrient-demanding as they need resources for growth and development (Dussutour and Simpson, 2008). We initially hypothesized that larvae, compared to workers, would show a similar or increased mortality as they get fed larger amounts of the test solutions and may be more susceptible. However, this was not the case for imidacloprid concentrations higher than 5 mg/L for which larval mortality was reduced compared to workers after 10 days. However, one reason for this is the high worker mortality (> 50%) itself. The death of workers caused impaired brood care leading to a reduced exposure of larvae to contaminated food. Larvae can survive without food for some time, meaning they were certainly starving. However, mortality caused by imidacloprid was lower due to reduced exposure, and starvation-related mortality was likely delayed beyond the 21-day duration of our experiments. Hence, if a significant portion of the worker force perishes, the immediate impact on larvae can be delayed or mitigated, as less brood care and particularly feeding reduces the uptake of toxicants but larvae will eventually die from starvation. Since larvae cannot survive on their own, exposure groups of the three highest concentrations showing high worker mortality and aberrant behavior were terminated after 10 days of exposure and larvae remaining without or less than 20% of the initial worker number per sub-colony were counted as dead larvae.

Measuring the active substance, in our case imidacloprid, in larval and pupal tissue would allow us to differentiate between the effect of the toxicant and starvation due to an increased worker mortality. However, such analyses exceed standard regulatory testing requirements. From a risk assessment perspective, the distinction between direct toxicant uptake by larvae or reduced brood care due to worker dysfunction is of less relevance. In both cases, the ultimate outcome — larval mortality and adverse effects on colony fitness—remains the same.

Larval survival until pupation (pupation rate) was significantly reduced at 1.7 mg/L and the LC_50_ was determined to be 0.77 mg/L (Fig. 4B). However, only 46 % of the larvae in the controls survived until pupation. The high control mortality of larvae is likely due to handling stress and frequent monitoring of larvae and pupae throughout the experiment. Manual inspections of the artificial nesting chambers every other day caused defensive behavior in workers, leading to the attempt to relocate larvae to less exposed areas. Frequent movement during development may have been a significant stressor, contributing to increased mortality. Consequently, we assume frequent movement during development may have been a significant stressor, contributing to increased mortality. For future testing, we recommend reducing the handling time, for example by using nest boxes that allow counting brood without handling it manually.

For each pupa, the day of pupation was recorded. Over the course of our level-2 experiment with *C. maculatus* only two larvae out of 60, exposed to 1.7 mg/L imidacloprid, pupated. Because larvae of varying age (selection criteria: 3–5 mm in length) were used, the day of pupation does not correspond to larval developmental time and variability was expected to be high. In consequence, no significant difference in the mean day of pupation was found up to 0.5 mg/L. Measuring developmental time is a well investigated sublethal endpoint to test for adverse effects of toxicants (Bartling et al., 2024; Gandara et al., 2024; van Loon et al., 2022). Testing effects on the development (e.g. time, growth, gene expression), are practical, reliable and feasible sublethal endpoints, if selection criteria for the larvae are defined (Gregorc et al., 2018; van Loon et al., 2022). For example, developmental time for larvae can be assessed when only larvae of the same size and weight are used for exposure experiments. A simple approach would be to include freshly hatched larvae, as older nymphs may vary in size and weight depending on their destined caste.

### Pupal development

We observed abnormal metamorphosis of some larvae, namely the absence of the protective layer (naked pupae in Fig. 5C compared to a regular pupa with pupal cocoon in Fig. 5D) that is typically found in species belonging to the subfamily Formicinae. Such naked pupae were only observed in exposure groups (NOEC <0.05 mg/L), but the proportion of naked pupae showed no monotonous concentration response relationship and thus, no significant response model could be fitted (Fig. 5A). These naked pupae were non-viable and, when left in their sub-colonies, were consumed by workers within 24 hours. This suggests that workers can detect non-viable individuals and remove dysfunctional pupae, consuming them as valuable resources.

**Figure 5:**
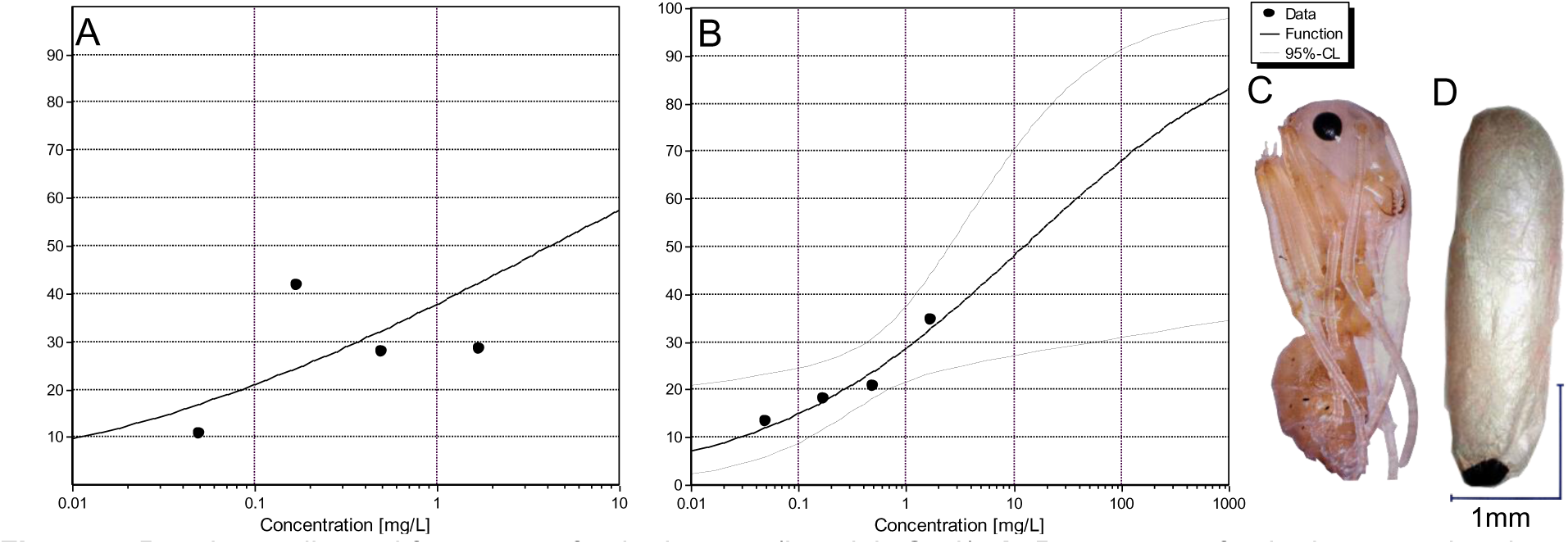
Pupal mortality and frequency of naked pupae (Level-2, Cm2): **A**: Percentage of naked pupae related to all pupae and **B**: Effect on pupal weight depending on imidacloprid concentration in food solutions. **C**: Naked pupae and **D**: Normal pupae with pupal cocoon

Additionally, the weight of viable pupae (with a pupal cocoon) was significantly reduced already at the lowest test concentration (NOEC < 0.05 mg/L) and the effect increased with increasing imidacloprid concentration in the food solutions (Fig. 5B). The EC_50_ was extrapolated to 18.1 mg/L.

Notably, naked pupae were only found in sub-colonies exposed to test solutions containing imidacloprid. Although the adverse and diverse effects on larvae could be demonstrated, the experimental design still needs further refinement to reduce larval mortality in the control sub-colonies.

### Summary

We demonstrated that workers, larvae and pupae strongly differ in sensitivity to imidacloprid. Pupal cocoon formation and pupal weight was affected already at the lowest test concentration where no effects of workers were observed. Abnormal larval development was influenced by direct effects of the toxicant and indirect effects due to reduced brood care of intoxicated and mortality of workers at higher concentrations. Since manually feeding ant larvae is impractical, measuring only the direct toxic effects on ant larvae is not feasible and thus indirect effects must always be considered. Our level-2 testing scheme can be further refined to assess additional sublethal effects, such as developmental time, brood-care frequency, and worker emergence rates (Fig. 1). We recommend focusing on pupation and emergence rates as reliable sublethal endpoints. However, these experiments require more time and labour compared to standard ecotoxicological tests. Since larval development and survival are critical for colony fitness, they should always be incorporated into the assessment of a toxicant’s hazardous potential.

#### Level-3 – Effects on queens and colonies

Newly mated *L. niger* queens were used as a proxy for the reproductive caste of entire colonies. *L. niger* queens start their colony independently and claustrally, i.e. individual founding queens’ do not leave their selected nesting place (subterranean or arboreal) and rely on their own reserves (fat body and flight muscles) to produce their first workers (Matte and Billen, 2021). In contrast to experiments with workers or larvae, individual founding queens were exposed only once to three different concentrations of imidacloprid at the beginning of the experiment (0.5, 5 and 50 mg/L, Fig. 2). This allowed us to investigate, whether single doses can have an adverse long-term effect on queens’ survival and reproductive behavior. Data on such feeding experiments are rare and to our knowledge were so far only conducted with semi-claustral queens of the Argentine ant (Rohel et al., 2001). Other studies tested the effects of pesticides on the survival and reproductive behavior on ant queens using topical application (Svoboda et al., 2023). However, since queens are almost always hidden in the safety of the nest (e.g. in the soil), topical application is not a field realistic scenario for ant queen pesticide exposure.

Survival of the founding queens over 21 days in the control group was high (95%). No significant effects on queen survival were observed up to 5 mg/L imidacloprid but mortality was significantly increased to 40 % at the highest tested concentration (Fig. 6A). Probit analysis calculated an LC_50_ of 111 mg/L. In contrast, our single exposure reduced queen reproduction already at the lowest test concentration (0.5 mg/L) significantly by 18 % (Fig. 6B). At a 100 times higher test concentration the effect was 40 % (Fig 6B). Since we used only three test concentrations the extrapolated concentration response (EC_50_ of 259 mg/L) was above the tested range and no confidence intervals could be calculated. Hence, this EC_50_ for *L. niger* needs to be confirmed by additional experiments.

**Figure 6:**
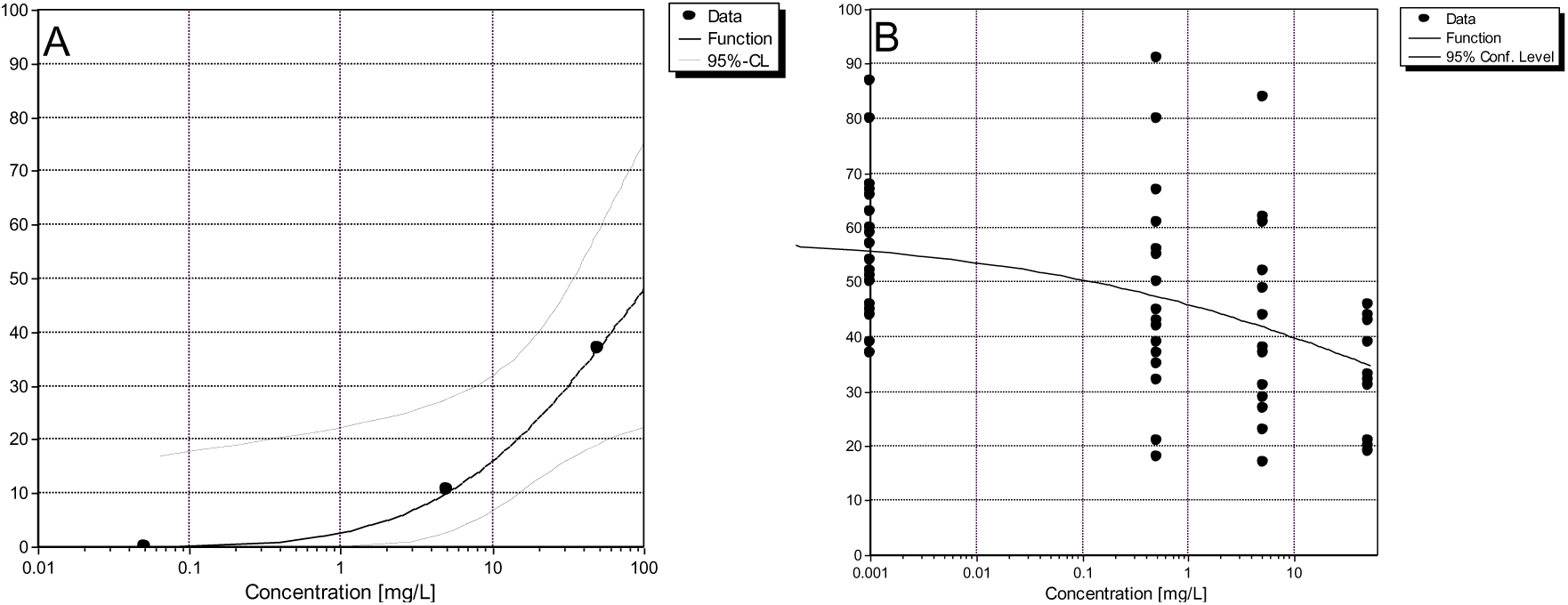
Survival and reproduction of queens over 3 weeks after a single dose at the beginning of the experiment (level-3). **A**: Probit analysis using Abbott’s formula to compensate for 5 % control mortality. **B**: 3-parameter log-logistic regression on the number eggs produced per surviving queen. The concentration in the control was set to 0.001 mg/L to allow log-transformation for fitting.

We consider using founding queens as a proxy for the reproductive caste of established colonies in a practical sense because they can be easily collected in the field and offer a cost- and time-effective alternative to testing entire colonies.

Testing whole colonies in level-3 is also an option which can be considered in future experiments. To reliably assess the impact of substances on different developmental stages of ant colonies, the effects of worker (or brood) loss should first be investigated, which is feasible but very time-consuming. One needs to collect for example 1200 founding queens (for *L. niger* and many other species, this can be done in one afternoon), bring them into the laboratory and set up for example three cohorts to test 400 founding colonies with 1-10 workers, 400 colonies with 50-100 workers (2^nd^ year) and 400 colonies with 500-1000 (3-4^th^ year) workers and perform systematic worker (and/or brood) removal experiments an determine the effect of this treatment on colony growth and survival. Using fast growing species without hibernation can be helpful to speed up this experiment. Once such work has been accomplished the data may be used to link experiments from level-1 and level-2 to colony growth or fitness.

### Summary

We have shown, that founding queens can serve as a proxy for the direct effects of imidacloprid on reproductively active queens (level-3). This can be a suitable option as fully grown colonies need months or even years to grow. An additional advantage of using founding queens is that one does not need to include indirect effects by workers in established colonies. We demonstrated that exposing claustral founding queens orally to pesticide is feasible. This experiment also demonstrated that only a single dose of imidacloprid at 0.5 mg/L has negative effects on the reproductive behavior of founding queens and may cause mortality at higher concentrations.

This result is significant in two ways when assessing the impact of chemical stressors on ant colonies. Experiments with founding queens not only help estimate effects on colony reproduction but also provide insights into the challenges associated with colony establishment. Colony founding is arguably the riskiest stage in an ant colonies life, e.g. Tschinkel and King (2017) showed in a large-scale field experiment that only 0.5 % founding queens of *Solenopsis invicta* survived the first 120 days. Additionally, observations of thousands of founding colonies over the last 30 years (Gadau personal observation) showed that those which lost their first set or workers never recovered from this event and eventually died. Therefore, any negative effect during this critical stage must have significant fitness effects for individual founding queens and in consequence for colony reproduction and finally population survival.

## Conclusions

### Imidacloprid as reference substance

Imidacloprid as a neonicotinoid is known to have delayed toxicity with accumulating effects over longer periods of time (Rondeau et al., 2014). Previous experiments using ants and bees already demonstrated that it influences behaviour, colony size and infection risks (Frizzi et al., 2023; Sappington, 2018; Schläppi et al., 2021, 2020; Wang et al., 2015), as well as the survival, foraging behavior, and (indirectly) reproduction (Bortolotti et al., 2003; Dively et al., 2015; Fischer, 2015; Parkinson and Gray, 2019; Ramirez-Romero et al., 2005; Yang et al., 2008).

Our experiments corroborated these findings and further demonstrated concentration dependent lethal and additional sublethal effects. Thus, imidacloprid is a good candidate for a reference substance (positive control) in further tests with ants.

### Caveats, problems and solutions

High larval mortality in our controls, which we observed in our initial experimental settings, can be avoided by reducing larval handling by the experimenter. To do so, we suggest assessing larval survival as the percentage of larvae that successfully pupated over a fixed amount of time. The use of founding queens as a proxy for colony level testing was successful but it can only estimate the direct effects of the stressor on queens (mortality and reproduction) and does not consider buffering or multiplication of effects due to queens being fed by workers in established colonies. Level-2 and level -3 experiments always should take indirect effects into account, e.g. since larvae and queens are highly dependent on workers for their survival or reproduction stressors may also act indirectly via reduced brood care which is also true for queens who once established do usually not actively forage. A potential solution, to disentangle direct from indirect effects, are either replacing exposed workers by control workers, that would only work reasonably well with one time exposure experiments or cross foster exposed larvae to control colonies.

### Advantages and Feasibility of the test system

The interest in ants so far has been mostly in terms of pest control due to the large number of invasive ants and their many negative effects (e.g. Siddiqui et al., 2021). However, although ants play many essential roles in most terrestrial ecosystems (Del Toro et al., 2012; Folgarait, 1998; Lobry De Bruyn, 1999), few studies have estimated the effects of PPP on ants as non-target organisms in ecotoxicological studies (Sappington, 2018; Schläppi et al., 2020; Thiel and Köhler, 2016; Wang et al., 2015) So far, ants are not part of a regulatory system to test for adverse effects on non-target organism.

By developing a formal test system and putting it into practice, we demonstrated, that ants can be efficiently used in ecotoxicity testing. In this context, ants have the advantage of easy handling and little requirements for space in the laboratory and therefore low maintenance costs, e.g. in comparison to honey bees or bumble bees. There is a long history and rich literature, both by professionals and amateurs, on how to rear hundreds of ant species from all over the world (https://www.antwiki.org/wiki/Welcome_to_AntWiki). The diversity and wide distribution of ants also has the advantage, that one can choose locally or globally occurring, ecologically specialized or generalist species (predators, herbivores, omnivores). Many ant colonies of various species can be kept for years under laboratory condition and many species (e.g. the three used here) show a high enough reproductive rate which allows to repeatedly establish multiple queenless sub-colonies (workers with or without brood) for a variety of experimental setups. This also has the advantage of performing tests on the same genotypes over many years increasing repeatability. However, we also showed species that require an obligatory diapause and exhibit medium growth rate can be successfully included in the testing scheme.

We also demonstrated that several organizational layers can be tested in ants with both lethal and sublethal parameters (Fig. 1 and Tab. 2). Ants may be especially suited for the detection of sublethal effects from stressors on non-target arthropods. Several reasons account for this: The complex eusocial structure of ants is sensitive to minor changes in brood care, communication or foraging which can be readily measured in the laboratory. Additionally, many individual (behavior, physiology, development) and colony level (growth, division of labour) fitness components can be determined in the laboratory on many experimental replicates in a reasonable time frame (weeks) and low cost. Compared to bees, the to-date only eusocial test organism in regulatory risk assessment, complete ant colonies can be kept and investigated in large numbers (100-1000) in the laboratory and therefore allow quantitative measures of sublethal effects and relate them to colony growth. Most ant species, even specialized predators like ponerines, feed on sugary solutions, if offered in the laboratory, which makes oral exposure and controlling uptake straightforward. Hence, a test system with ants is very versatile and can be tailored to specific applications.

Summarizing, we highly recommend using ants as new model organisms in future ecotoxicological testing. This study can be seen as starting point for refined testing.

### Linking hazard to colony fitness

Ecotoxicity testing is an important aspect to ensure that products such as biocides or pesticides, including plant integrated pesticides in GMO, are safe for animal health, and the environment. While toxicity of pesticides decreased over time for mammals, this is not necessarily the case for arthropods (Schulz et al. 2021). In particular, the massive loss of biodiversity over the last decades in agriculturally dominated landscapes makes it necessary to reduce effects on non-target arthropods. Coming back to our introduction, we see many advantages including ants as non-target test organisms in ecotoxicity testing. Especially to protect biodiversity, for which new testing methods have been stipulated by the EU authorities (EFSA 2015). To do so the versatile test system needs to be standardized and linked to a critical damage threshold in the sense of a specific protection goal (SPG) for ants (EFSA, 2016). Therefore, we propose to link potential parameters (Fig. 2) to colony fitness and to define a harmful effect on colonies in terms of colony growth or fitness which can act as specific protection goal. It is straight forward to estimate direct and even indirect measures of lethal and sublethal effects on individual larvae, workers or queens but how these effects scale to the colony level is not known in detail and may also depend on the species tested. Many of the suggested parameters as endpoints here (worker survival, larval survival, reproduction of the queen), link to colony growth. Negative growth itself leads to less workers which lead to a reduction of brood and eventually to a downward spiral ultimately ending in colony death. Also, in single queen colonies the death of the queen results in the death of her colony. Questions on how much additional worker mortality or how many changes in worker behavior can be buffered by a colony cannot be answered quantitatively. Nobody has so far done an experimental test removing an increasing amount of the workforce to empirically demonstrate a threshold for an acceptable loss of the workforce that would allow a colony to recover. Such a threshold is certainly species specific but also depends on the developmental stage of a colony. Founding colonies with few workers are much more susceptible to worker loss than large and established colonies which regularly loose thousands of workers (Kwapich and Tschinkel, 2013).

In general, the evaluation of a single sublethal effects may not be suitable to predict colony effects, as the overall effect on the colony fitness is driven by a combination of both lethal and sublethal effects on every caste (workers, brood, queen, alates). To evaluate the impact on colony fitness, developing a theoretical model that integrates sublethal and lethal effects would be highly useful to define damage scenarios and thresholds for adverse effects.

## Acknowledgement

This publication relates to the Research & Development Project (F+E) No 3520 84 0400 commissioned by the Federal Agency for Nature Conservation Germany with funds from the Federal Ministry for the Environment, Nature Conservation, Nuclear Safety and Consumer Protection.

## Authorship contribution

MO and JG conceived of the study; MP and MO drafted the manuscript; MP maintained the colonies and carried out experiments for the case study. BR and BG were responsible for the chemical analysis of imidacloprid. UH, SE, CS and contributed to the revision of the manuscript and to the discussion and approved the final version.

## Notes

### Competing Interest Statement

The authors have declared no competing interest.

